# Photocatalytic disinfection of Surfaces with Copper Doped TiO_2_ Nanotube Coatings Illuminated by Ceiling Mounted Fluorescent Light

**DOI:** 10.1101/225615

**Authors:** Tilen Koklic, Štefan Pintarič, Irena Zdovc, Majda Golob, Polona Umek, Alma Mehle, Martin Dobeic, Janez Štrancar

**Author notes:** contributed equally to this work (Tilen Koklic, Štefan Pintarič).

## Abstract

High economic burden is associated with foodborne illnesses. Different disinfection methods are therefore employed in food processing industry; such as use of ultraviolet light or usage of surfaces with copper-containing alloys. However, all the disinfection methods currently in use have some shortcomings. Here we show that copper doped TiO_2_ nanotubes deposited on existing surfaces and illuminated with ceiling mounted fluorescent lights or additional low power light emitting diodes can be employed for an economical and permanent disinfection of surfaces.

We deposited the nanotubes on various surfaces: polyethylene terephatlate, polystyrene, and aluminum oxide, where they could withstand repeated washings with neutral, alkaline or acidic medium. Here we show that the polymer surfaces coated with the nanotubes and innoculated with 10^7^ bacteria, illuminated with ceiling mounted fluorescent lights retard the growth of *Listeria Innocua* by up to 99% in seven hours of exposure to the fluorescent lights, compared to a control surface. The disinfection properties of the surfaces depend mainly on the temperature difference of the surface and the dew point, where for maximum effectiveness of the photocatalytic effect the difference should be at least 2.5 degrees celsius.

Usage of one dimensional nanomaterials, such as TiO_2_ nanotubes, offers a promising low cost alternative to current disinfection methods, since illumination of surfaces with common fluorescent lights is sufficient to photo-excite the nanotubes, which sequentially produce microbicidal hydroxyl radicals. Future use of such surfaces with antibacterial nano-coating and resulting sterilizing effect holds promise for such materials to be used in different environments or in better control of critical control points in food production as well as an improved biosecurity during the food manufacturing process.

## Introduction

Economic burden of $30–80 billion was estimated by the Center for Disease Control and Prevention (CDC) for the annual number of foodborne illnesses, affecting 48 million Americans ^1,2^. Over 320.000 cases of food-borne zoonotic diseases were evidenced in humans each year, thus the measures in view of food safety have to be very strict especially on food and food premises hygiene ^3^. Food can become contaminated at any point during production and distribution, as well as in consumers’ own kitchens. Therefore, foodborne illness risk reduction and control interventions must be implemented at every step throughout the food preparation process ^4^. Recent global developments are increasingly challenging international health security according to the World Health Organization (WHO). These developments include the growing industrialization and trade of food production and the emergence of new or antibiotic-resistant pathogens. Micro-organisms are known to survive on surfaces, for extended periods of time. Among the foodborne pathogens, *Listeria monocytogenes* has the highest mortality rate in humans and is one of the most environmentally resistant facultative anaerobic bacteria growing at its optimal temperatures from −18 °C to 10 °C, in environments with or without oxygen with propensity of forming a biofilm ^5^. Between 13 serotypes of *Listeria monocytogenes*, three serotypes (1/2a, 1/2b and 4b) are the reason for the majority of human listeriosis ^6^. In our previous research using pulsed-field gel electrophoresis typing of *L. monocytogenes* isolates from poultry abattoir we identified the same serotype (classical 1/2a, molecular IIa) with the exception of one isolate with a different serotype (4b, IVb), mainly found on the surface, but some also in the air ^7^.

Many disinfectants were tested in the prevention of *Listeria monocytogenes* contamination, however organic burdening and biofilm formation effectively inhibited disinfectants’ microbicidal activity ^8,9^. Although biofilm formation is common for every environment where microorganisms are close to the surface, its formation is even more problematic in the food industry, where remains of foods in inaccessible places enable survival and the multiplication of *Listeria*. It was speculated that specific properties of persistence of *L. monocytogenes*, might be the reason for spreading of persistent strains of *L. monocytogenes* across the surfaces of food-processing plants, but also by transferring meat products between different plants ^10,11^. In addition, some studies report about the possibility of reduced *L. monocytogenes* susceptibility to some chemical disinfectants ^12^. Permanent maintenance of hygiene in food processing industry is therefore of utmost importance for the continuous reduction in the number of bacteria. For this reason, regular cleaning and disinfection is mandatory, but it is often performed poorly and irregularly specially when parts of the meat processing equipment are inaccessible ^13^. Namely the risk for food contamination arise mainly due to low hygiene of food premises and not from previously contaminated animals as it was shown by Ojeniyi et al. ^14^, and by our own work, where we were unable to confirm the transfer of *L monocytogenes* from broiler farm to the abattoir, since we couldn’t prove a positive case of *L monocytogenes* on broiler farms among the investigated animals ^7^. One of the main reasons for spreading of the persistent strains of *L. monocytogenes* might be its ability of enhanced adherence to surfaces in a relatively short time ^15,16^, therefore the continuous antibacterial function of food contact surfaces should be implemented. One of such continuous disinfection methods, suitable for disinfection of the air, liquids and surfaces is the use of ultraviolet light (UV), which is being employed as one of the physical methods of decontamination in the food processing industry ^17^. Short-wave ultraviolet light (UVC, 254 nm) was shown to be effective against wide spectrum of bacteria, viruses, protozoa, fungi, yeasts and algae, by altering cell DNA ^17^. However, UVC has limited applicability in food industry since it can cause sunburn, skin cancer, and eye damage under direct exposure. UVC lights can also produce ozone, which can be harmful to human health, and finally materials exposed to UVC light for longer period age faster, especially plastics and rubber, which break down under UVC exposure. On the other hand, long-wave ultraviolet light (UVA, >320 nm) as a part of a sunlight, not absorbed by the atmosphere ozone layer, thus reaching the earth’s ground, and is not harmful to human health, can still cause some oxidative damage, however has much weaker effect on microorganisms than UVC ^17^. Since UVA is normally present as a small part of the fluorescent lighting spectrum, one could use ceiling mounted fluorescent lights for permanent surface disinfection provided that the oxidative damage of UVA light at a surface could be enhanced. This can be achieved by illuminating TiO_2_ deposited on a surface by UVA light. Namely, illuminated TiO_2_ is known to produce reactive oxygen species, such as hydroxyl or superoxide radicals, which can also be used for disinfection of surfaces. As early as in 1977 it has been shown that TiO_2_ can decompose cyanide in water when illuminated with sunlight ^18^. If TiO_2_ is irradiated with photons with energies greater than material’s band gap, electron-hole pairs are generated – for TiO_2_ with Eg around 3 eV wavelengths below approximately 415 nm are needed ^19^. Photo generated holes are highly oxidizing whereas photo generated electrons are reducing enough to produce superoxide from dioxygen ^20^. After reacting with water, holes can produce hydroxyl radicals (·OH). Photo excited electrons can become trapped and loose some of their reducing power, but are still capable of reducing dioxygen to superoxide radical (·O_2_-), or to hydrogen peroxide H_2_O_2_. Hydroxyl radical, superoxide radical, hydrogen peroxide, and molecular oxygen could all play important roles in preventing proliferation of bacteria. Using TiO_2_ surface coatings one should therefore be able to maintain clean surfaces with the use of UV light close to visible spectrum.

We have shown previously that Cu^2+^-doped TiO_2_ nanotubes (Cu-TiO_2_NTs) coated polymer surfaces reduce number of seeded bacteria by 99.94% ± 0.05% (i.e. Log_10_ reduction = 3.5 ± 0.5, when innoculated with 2.4 10^4^ *Listeria innocua*) when illuminated with low power UVA diodes for 24 hours at 4 °C ^21^, where the intensity of UVA light needed to observe the antibacterial effect was only about 10 times more than it is usually present in common fluorescent lighting. In this manuscript we present antibacterial effect observed on polymer surfaces when illuminated with common fluorescent lights, which were already present on a ceiling of a food processing plant. Coated surfaces innoculated with 10^7^ bacteria exhibit similar antimicrobial effect as we observed previously on the TiO_2_ nanotube coated petri dishes, reducing the number of *Listeria innocua* up to 99 *%* in seven hours of exposure to the fluorescent lights, compared to control surfaces.

## Materials and Methods

### Materials

The spin trap, 5-(Diethoxyphosphoryl)-5-methyl-1-pyrroline-N-oxide (DEPMPO) (Alexis, Lausen) was used as purchased without further purification and stored at −80 °C. The spin-trap stock solutions were always freshly prepared. Ethanol (EtOH) and methanol (MeOl·l) from Merck AG (Darmstadt, Germany) were used in Lichrosolv^®^ gradient grade quality. Media and culture materials were obtained from Gibco – Invitrogen Corporation (Carlsbad, California).

### Preparation of bacterial inoculum

Antimicrobial properties were tested on non-pathogenic bacterium *Listeria innocua*, which is closely related to pathogenic species *Listeria monocytogenes*. Suspension of *Listeria innocua* strain, isolated during routine examination (RDK.), was supplied by the Institute of Microbiology and Parasitology, Veterinary faculty, University of Ljubljana. Strain was maintained frozen at −70 °C in sterile vials containing porous beads which serve as carriers to support microorganisms (Microbank, pro-lab Diagnostics) and kept at −70 °C. The inoculum was prepared in liquid medium and incubated aerobically for 24 h at 37 °C. After incubation the culture contain approximately 10^9^ colony forming units (CFU) per milliliter. Working suspensions with appropriate concentrations were achieved by several 10-fold dilutions.

### Preparation and properties of Cu^2+^-doped TiO_2_ nanotubes

Cu^2+^-doped TiO_2_ nanotubes (Cu-TiO_2_NTs) were prepared in several steps: (i) first sodium titanate nanotubes (NaTiNTs) were synthesized from anatase powder (325 mesh, ≥ 99.9%, Aldrich) and 10 M NaOH (aq) (Aldrich) at T = 135 °C for 3 days under hydrothermal conditions. Exact synthesis procedure is described previously ^22^, (ii) in the next step NaTiNTs were rinsed with 0.1 M HCl(aq) yielding protonated titanate nanotubes (HTiNTs), (iii) then 400 mg HTiNTs were dispersed in 100 mL of 0.5 mM solution of Cu^2+^(aq) (source of the Cu^2+^ was CuSO_4_·5H_2_O (Riedel de Haen)) using an ultrasonic bath (30 minutes) and stirred at room temperature for 3 hours. By centrifugation the solid material was separated from the solution, and (iv) finally isolated material was heated in air at 375 °C for 10 hours.

The powder X-ray diffraction (XRD) pattern was obtained on a Bruker AXS D4 Endeavor diffractometer using Cu Kα radiation (1.5406 Å; in the 2*θ* range from 10 to 65°). Morphology of the particles in the sample was determined using transmission electron microscope (TEM, Jeol 2100). The specimen for the TEM investigation was prepared by dispersing the sample in MeOH with the help of an ultrasonic bath and depositing a droplet of the dispersion on a lacey carbon-coated copper grid.

### Activity of TiO_2_ nanotubes

The photocatalytic activity of synthesized titanate and TiO_2_ nanomaterials was determined using electron paramagnetic resonance spectroscopy (EPR) with spin trapping, which was optimized for measurement of primary radicals generated in the vicinity of the nanomaterial surface. This was achieved by measuring primary hidroxyl radicals in the presence of 30% ethanol with 5-(diethoxyphosphoryl)-5-methyl-1-pyrroline-N-oxide spin trap (DEPMPO). EPR spin trapping was applied to measure the generation of reactive oxygen species (ROS) production.

### Deposition of Cu-TiO_2_NTs on PET surface and testing of the deposition stability

The deposition of Cu-TiO_2_NTs was made on different surfaces: 2.5 cm × 7.5 cm polyethylene terephthalate (PET) slides, polystyrene petri dishes (8 cm diameter), aluminum oxide slides with the same dimensions as PET slides. The surfaces were washed before deposition. They were soaked in 20% NaOH solution, rinsed with distilled water, and finally with ethanol vapor.

The suspension of the nanotubes with concentration of 1 mg/mL was processed with ultrasonic liquid processor (Sonicator 4000, Misonix) prior to the deposition on the surfaces. Sonication was performed using 419 Microtip™ probe, 15 min process time, 10 s pulse-ON time, 10 s pulse-OFF time and maximum amplitude (resulting in 52 W of power).

The surfaces were treated with compressed air 3 times for 3 s. 150 μL of nanoparticle suspension was applied on each surface, immediately after compressed air treatment, and smeared evenly. The same number of surfaces with nanoparticle deposition and control surfaces were prepared for each experiment. On control surfaces, only 150 μL of solution was applied. After the deposition, the surfaces were left in the oven at 50 °C for 2 h. Then they were rinsed with distilled water and put back in the oven at 50 °C for another 2 h.

The photocatalytic activity of the nanodeposit on the surfaces was tested using EPR spectroscopy. Three measurements were performed on the surfaces, with or without the nanodeposit. On each surface, small pool, proportionate to the size of the sample, was made with silicon paste and was filled with 2 μL of 0,5 M DEPMPO and 18 μl of 30% ethanol and irradiated with 290 nm diode for 5 min. The diode was 1–2 mm above the surface of the sample. The solution with short-lived radicals being trapped in the form of stable DEPMPO spin adducts was then drawn into the quartz capillary of 1 mm diameter, which was put in the 5 mm wide quartz tube and transferred into EPR spectrometer. All EPR measurements were performed on an X-band EPR spectrometer Bruker ELEXYS, Type W3002180. All measurements were recorded at room temperature using 1 Gauss (10^-4^ T) modulation amplitude, 100 kHz modulation frequency, 20 ms time constant, 15 x 20 seconds sweep time, 20 mW microwave power and 150 G sweep width with center field positioned at 3320 G.

The amount of deposited material was estimated from EPR signal decrease when rinsing the deposit of 150 μL of 1 mg/mL applied to a 2.5 × 7.5=18.8 cm^2^ surface. With EPR signal being decreased to about 1/3, we estimated that the amount of deposited nanomaterial was about 2 μg/cm^2^.

### Antimicrobial Activity of nanotube coated PET surface in a meat processing plant

Four measurement points were selected in a poultry slaughterhouse with regards to different air microclimate conditions (humidity, temperature, airflow) as well as intensity of UV irradiation. PET slides were innoculated with 10^7^ bacteria in 10 μL droplet and placed either vertically or horizontally at different altitudes (0.5 or 2 m) and exposed for 7 hours. After exposure, the samples were washed in saline (NaCl 0.9%) and examined bacteriologically to determine the number of bacteria. Survival of bacterial culture of *Listeria innocua* has been measured for samples with and without germicidal Cu-TiO_2_NTs coating. Reduction ratio was expressed in percentage and logarithm (Log_10_). % reduction was calculated using the following equation:

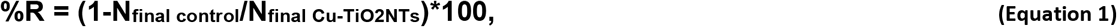

Where N_final control_ is the number of bacteria after the exposure on a control surface, and N_final Cu-TiO2NTs_ is the number of bacteria after the exposure on a surface coated with Cu-TiO_2_NTs nanotubes.

Log reduction was calculated using the following equation:

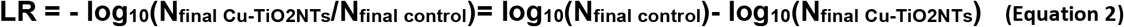

### Antimicrobial activity in presence of repeated daily contamination and washing

Effect of long term illumination was studied only on PET surfaces, by placing uncoated (control) PET slides and PET slides covered by Cu-TiO_2_NTs on cooled (4 °C) aluminum plates in order to mimic cold and condensing conditions at the cooling walls commonly present in food processing plants. Bacterial suspension (10 μl) of living microorganism *Listeria innocua* in concentration of 1.5 to 5.0 × 10^9^ CFU/mL was applied daily on each PET slide. The slides were then cooled to the dew point, which prevented the drying of microorganism containing droplets on the slides. Slides were washed with 100 mL of sterile saline solution (0.9 weigth % NaCl) at different time intervals and the number of surviving microorganisms was determined. The remaining PET slides were stored in the dark at 4 °C until the next day when the above described process was repeated. The whole experiment with daily washing and bacteria application lasted for 28 days.

## Results and Discussion

### Structure and photochemical activity of copper doped TiO_2_ nanotubes (Cu-TiO_2_NTs)

Transmission electron microscopy (TEM) images (*Figure 1* A and B) show that nanotube morphology is maintained after incorporation of copper ions, albeit images taken at higher magnifications *Figure 1* B reveal that nanotube walls are not as clearly defined as in original sodium titanate nanotubes, described previously ^23^. More detailed characterization, using also advanced high resolutiom transmission electron microscopy techniques, of the copper doped TiO_2_ nanotubes is reported in Koklic et al. ^21^ (article submitted, please see the Supporting Information for Review Only). X-ray diffractogram (XRD) of the Cu-TiO_2_NTs nanotubes is shown in *Figure 1* C, where all peaks correspond to anatase TiO_2_ (JCPDS No. 89–4203). Elemental analysis (EDS) indicated (data not shown) that the copper content is about 0.1 weight %.

**Figure 1.**
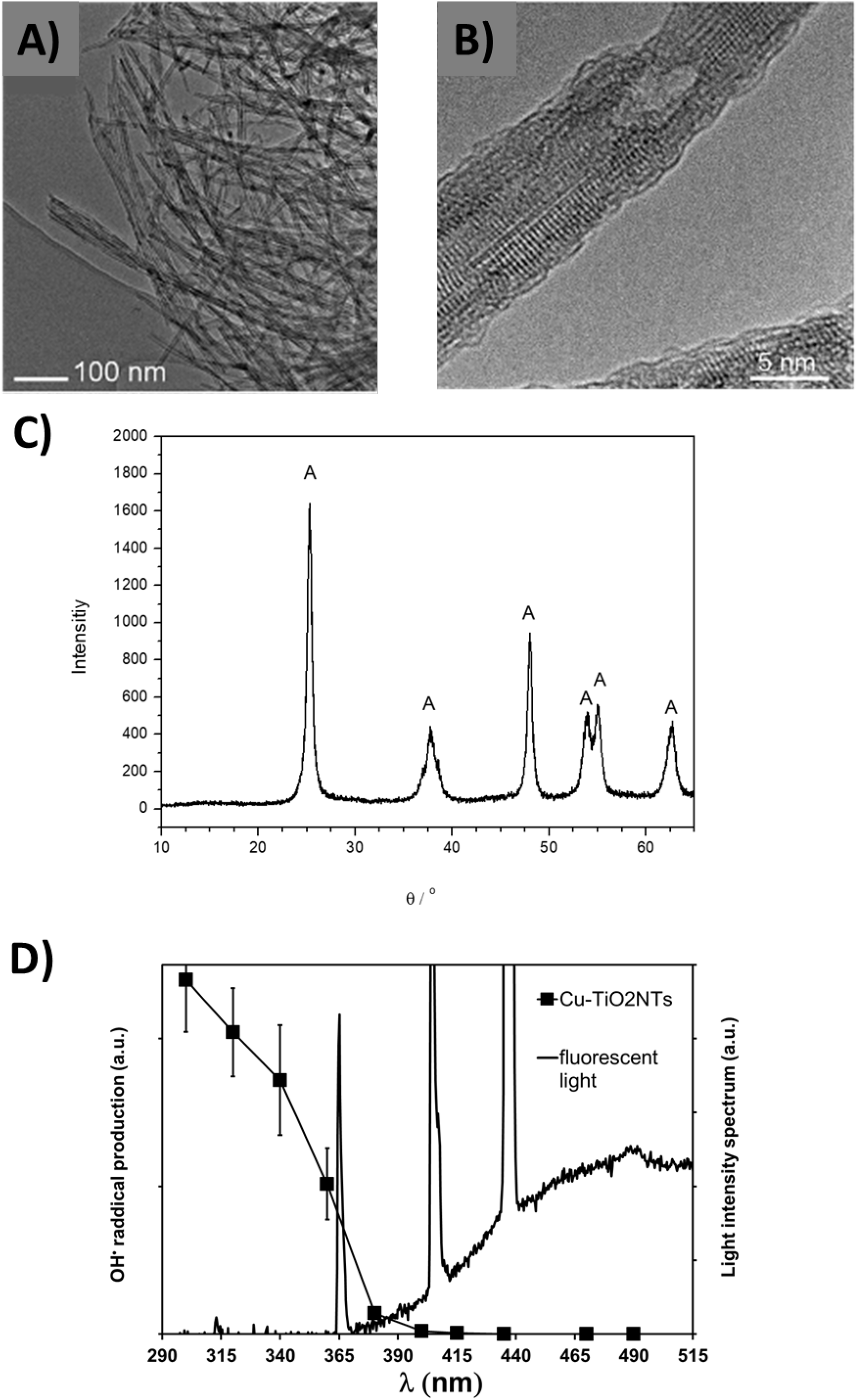
Structure and photocatalytic activity of Cu^2+^-doped TiO_2_ nanotubes (Cu-TiO_2_NTs) excited at different wavelengths. **A)** and **B)** TEM images of the nanotubes taken at different magnifications; **C)** XRD of the nanotubes. Anatase peaks are marked with A; **D)** Amount of hydroxyl radical production (closed squares) versus illumination of Cu-TiO_2_NTs at different wavelengths is shown against emitted light spectrum of a common fluorescent light (black line).

We have previously shown that the Cu-TiO_2_NTs deposited on polystyrene petri dishes reduce up to 99.94% ± 0.05 (3.5±0.05 log10 reduction, initial number of bacteria 2.5·10^4^) *Listeria innocua* in 24 hours in a refrigerator at 100% humidity, illuminated with UVA light emitting diodes ^21^. However, Usage of additional illumination results in additional costs associated with application of such disinfection methods. On the other hand, ceiling mounted fluorescent lights, which are already in use in many food processing plants, contain a small portion of emitted light in UVA range. *Figure 1* D shows the spectrum of emitted light by a ceiling mounted fluorescent lamp. Three peaks in the spectrum are clearly visible, with one spectral peak at 365 nm. It is this peak which is absorbed by the Cu-TiO_2_NTs, as it is evident from the absorption of light as a function wavelength (*Figure 1* D, closed squares). The absorption of light versus wavelength of the light is consistent with a bandgap of the TiO_2_, a property of a semiconductor such as TiO_2_. Matsunaga et al. showed already in 1985 that Escherichia coli cells were completely sterilized when TiO_2_ was irradiated with UV light.^24^ Since then the antibacterial effect of photoexcited TiO_2_ was shown against a wide range of microorganisms.^25^ Photocatalytic mechanism and related photochemistry of TiO_2_ is well researched^26–29,29–33^, antibacterial action seems to depend mainly on ·OH radicals, which are produced on the surface of TiO_2_ when illuminated with light consisting of wavelengths below TiO_2_’s bandgap. Due to this semiconductor property of the Cu-TiO_2_NTs nanotubes the production of hydroxyl (OH·) radicals on the surface of nanotubes increases with decreasing wavelength. We measured the amount of produced radicals as a function of different wavelengths of light, by using a DEPMPO spin trap (*Figure 1* D, closed squares), which is commonly used for efficient trapping of the hydroxyl radicals ^34^. Since the spectrum of the emitted light from a ceiling mounted common fluorescent light (*Figure 1* D, black line, the peak at 365 nm) overlaps with the spectrum of the light needed for efficient photoexcitation of the nanotubes (the closed squares), we expected that the nanotube coated surfaces could be excited by fluorescent lights, which are already present on ceilings at food processing plants. Especially due to intense peak at 365 nm, which is present in the emitted spectrum of the fluorescent light bulb and represents about 1% of total light emitted by the lamp.

### Deposition stability of copper doped TiO_2_ nanotubes on different surfaces

Next we tested the stability of the nanotubes deposited on different surfaces, which are commonly used in food processing industry. The dispersion of Cu-TiO_2_NTs was added to the clean surface (see Materials and methods) and left to dry. No special chemical modification of either nanotubes or surface was necessary. Unattached nanoparticles were washed away under a stream of water and the production of hydroxyl radicals was measured as described in Materials and Methods section. Since the amount of the produced radicals is proportional to the amount of Cu-TiO_2_NTs still remaining on the surface after extensive washing, we used the measurement of the the quantity of radicals produced by illuinated surfaces as a measure for the stability of the deposition. That is, if the amount of produced radicals remains constant throughout the washing cycles, then the Cu-TiO_2_NTs nanoparticles should remain attached to the surface. We tested different materials: polyethylene terephtalate (Figure 2, row A), polystirene (Figure 2, row B), and Aluminum oxide (Figure 2, row C). All surfaces were repeatedly soaked at different pH conditions neutral (pH7, Figure 2, first column, blue color), basic (pH10, Figure 2, second column, violet color), and acidic (pH4, Figure 2, third column, red color) and extensively washed under a stream of water after each soaking. The amount of material versus washing step was fit with a linear curve using GraphPad Prism version 7.00 for Windows (GraphPad Software, La Jolla California USA, www.graphpad.com). The area, which contains a linear fit, which describes the data with 90% certainty is shown on each graph. In all of the graphs, except for aluminum oxide washed at pH10, linear fit is contained around 100% deposited material (horizontal dotted lines), thus indicating that Cu-TiO_2_NTs nanoparticles deposited to various surfaces should withstand daily washings with different detergents commonly used in food processing industry. However the material will not provide long term disinfection of aluminum oxide surfaces, when washed with basic detergent (Figure 2, frame C2).

**Figure 2.**
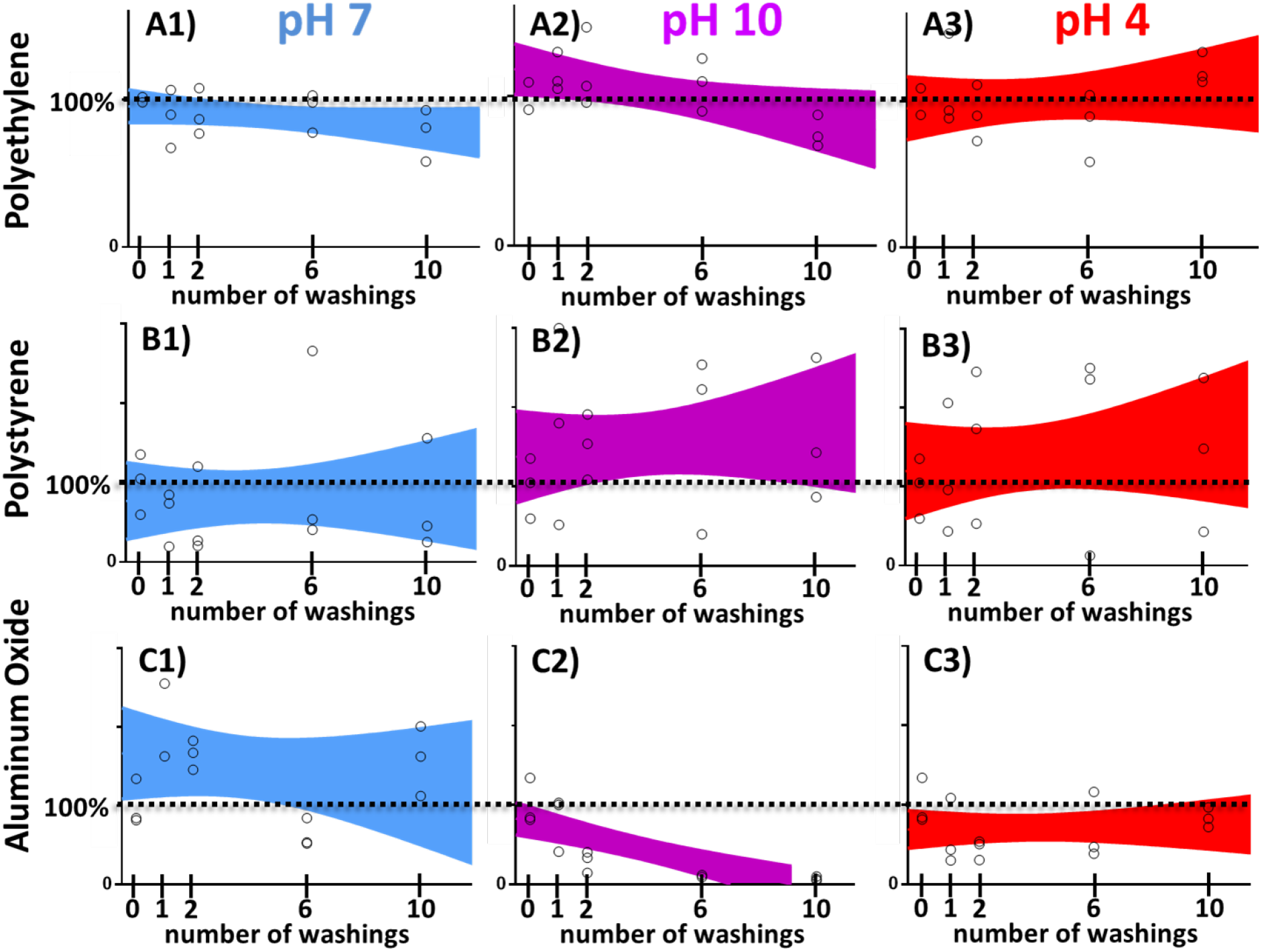
Deposition stability of copper doped TiO_2_ nanotube coatings on different surfaces. Stability on the surface against washing at different pH conditions (neutral, pH7 - blue, acidic, pH4 – red, basic, pH10 – violet), without abrasion, but under extensive water flow. Individual measurements are shown as open circles; colored areas represent 90% confidence areas, which enclose the area that one can be 90% sure to contain the linear fit curve.

### Antimicrobial activity of TiO_2_ nanotube coated surfaces placed in a food processing plant

We exposed polyethylene terephthalate (PET) surfaces with or without Cu-TiO_2_NTs antibacterial coating at different locations in the food processing plant to test whether the intensity of ceiling mounted fluorescent lights in a food processing plant is sufficient to provide measurable antibacterial activity of surfaces coated with the nanotubes. We applied 10 μL of bacterial suspension of *Listeria innocua* on the PET surfaces (10^7^ bacteria), as shown schematically in Figure 2 A, and placed the PET surfaces at different places in the food processing plant with respect to performed tasks (evisceration, meat cut up, cold room, Depo – meat storage, and butchering) for 7 hours. The air microclimate conditions (UV light Intensity, humidity, and temperature) were followed and the number of remaining bacteria was determined (Table in Figure 3 C). Microclimatic air conditions measured at different places in the food processing plant were different with respect to temperature, humidity, ambient light intensity emitted from ceiling mounted fluorescent lights, and airflow (Figure S 1). We measured the highest disinfection activity in meat cut up room, where the reduction of the number of *Listeria Innocua* was 99% in seven hours of exposure to the fluorescent lights, compared to a control surface. The reduction of the number of bacteria was high at three places: 1) Evisceration (90%), 2) Cut up (99%), and 3) Cold room (73%) (*Figure 3* D, grey bars). The disinfection properties of the surfaces depend mainly on the temperature difference of the surface and the dew point (*Figure 3* D, black bars), where for maximum effectiveness of the photocatalytic effect the difference should be less than about 2.5 °C (*Figure 3* D, black dashed line). This is not supprising, since fogs of all types start forming when the air temperature and dewpoint of the air become nearly identical. This occurs through cooling of the air near the cool surface to a little beyond its dewpoint and the precipitation of water droplet from the air seldom forms when the dewpoint spread is greater than 2.5 °C. Other microclimatic parameters (Figure S1) didn’t follow the relative reduction of *L. innocua*. This results shows that photocatalytic disinfection of surfaces can be made efficient in humid places of the food processing plant. On the other hand, significant reduction of the number of bacteria was also found in the number of remaining bacteria on nanotube coated surfaces versus uncoated control surfaces (*Figure 3* E) when we averaged reductions of bacteria across all the places in the food processing plant. The average number of bacteria on the surfaces with the nanotube coating was reduced more (Log10 = 2.92) than on the control surfaces (Log_10_ = 3.90), which can be expressed as relative percent reduction of %R = 90%.

**Figure 3.**
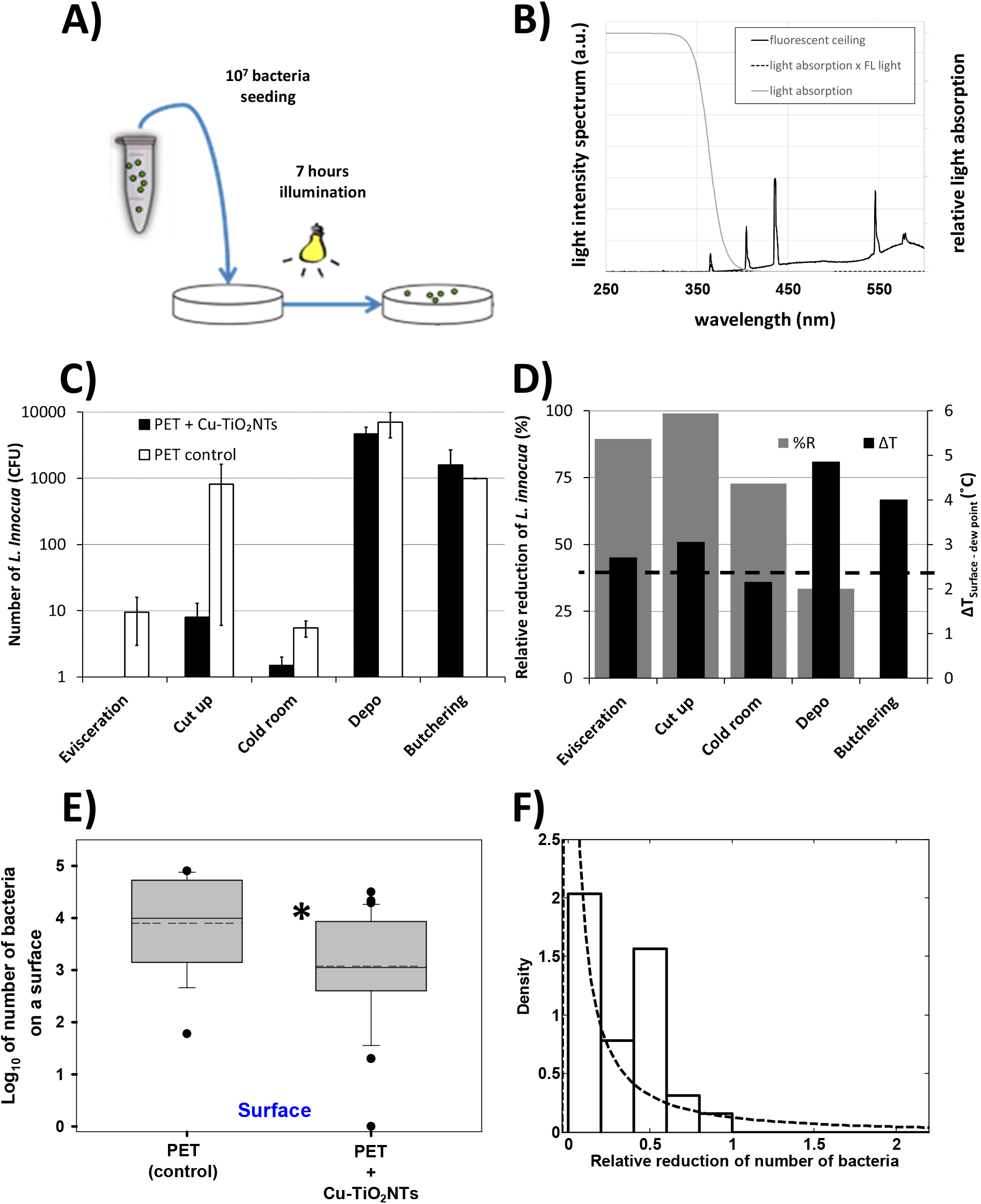
Survival of Listeria innocua under exposure to ceiling mounted fluorescent lights in a food processing plant. A) Schematic presentation of the experiment. B) spectrum of emited light from ceiling mounted fluorescent lights (black line); relative light absorption by the Cu-TiO_2_NTs (grey line); spectrum of absorbed light by the nanotubes (dashed line). C) Number of Listeria innocua shown on a logarithmic scale. On each surface 10^7^ bacteria were placed and left at different places in the food processing plant for 7 hours. After this time period remaining bacteria were transferred from the surfaces and colony forming units (CFU) were counted. Number of CFU on control surfaces is shown with white bars (PET control); Number of CFU on the nanoparticle coated surfaces is shown with black bars (PET + Cu-TiO_2_NTs); D) Relative reduction (%R) of bacteria as a consequnce of disinfecting action of nanoparticle coated surface, illuminated with ceiling mounted fluorescent lights (grey bars), calculated according to the equation 1 in Materials and Methods section; Black bars represent the microclimatic parameter (difference between surface temperature and dew point temperature – dew point spread) which correlates with the %R. E) Boxplot presents log10 of number of bacteria on either control PET surface without antibacterial nano coating (PET) or the number of bacteria on a PET surface coated with copper doped TiO_2_ nanotubes (PET+Cu-TiO_2_NTs). Median value of each distribution is shown with horizontal line within each box, while the dashed line marks the mean. The boundary of the box closest to zero indicates 25th percentile, and the boundary farthest from zero indicates the 75th percentile. Whiskers (error bars) above and below the box indicate the 90th and 10th percentiles. The outliers are shown as dots. Note the LOG scale. Since normality test (Shapiro-Wilk) failed (P < 0.050), Mann-Whitney rank sum test *was* performed on the control group (N=16, median=4) and on the PET+Cu-TiO_2_NTs group (N=32, median=3.1). The difference in the median values between the two groups is greater than would be expected by chance (P = 0.003)^*^. F) Histogram of the distribution (probability density function – PDF) of survival ratios, calculated as a ratio between the number of Listeria innocua from the surface with antibacterial coating (PET+Cu-TiO_2_NTs) and the number of Listeria innocua from a control surface without the coating (PET). For inefficient antibacterial coating survival ratio of 1 is expected. Dotted line is lognormal fit of the distribution with a maximum of the PDF below 0.1 for the reduction of the number of bacteria on the surface.

For further quantification of the results we calculated the ratios of bacteria from the coated surfaces versus the control surfaces for all the measurements (*Figure 3* D, white bars). In such presentation of the results the antibacterial effect is reflected in the ratio to of less than 1. The distributions of survival ratios (*Figure 3* D, white bars) were clearly not normal. Since biological mechanisms often induce lognormal distributions ^35^, for example when exponential growth is combined with further symmetrical variation such as initial concentration of bacteria ^36–39^, we fit our data with a log normal distribution (*Figure 3* D, dashed line). The lognormal fits of the histograms fit best the survival ratios also when compared to other distributions.

### Antimicrobial activity in presence of repeated daily contamination and washing

Next we repeatedly inoculated and washed PET surfaces with *Listeria innocua* daily, in order to mimic daily contamination in food processing industry or surfaces in a household, as shown schematically in *Figure 4* A. After application, we left the bacteria on the surface for 7 hours while being exposed to low intensity light from fluorescent lamps on the ceiling (t=7 h, j=2.5 W/m^2^, A= 8 J (total light), A_<380nm_= 80 mJ). As it can be seen from the measured light intensity spectrum of the fluorescent lamp (*Figure 4* B), the intensity of light with wavelengths below 380 nm (80 mJ) is only 1 percent of the total light intensity, the corresponding energy of the illumination of the nanotubes, which can induce the photocatalytic process of hydroxyl radical production, is therefore around 80 mJ in our experimental setup.

**Figure 4.**
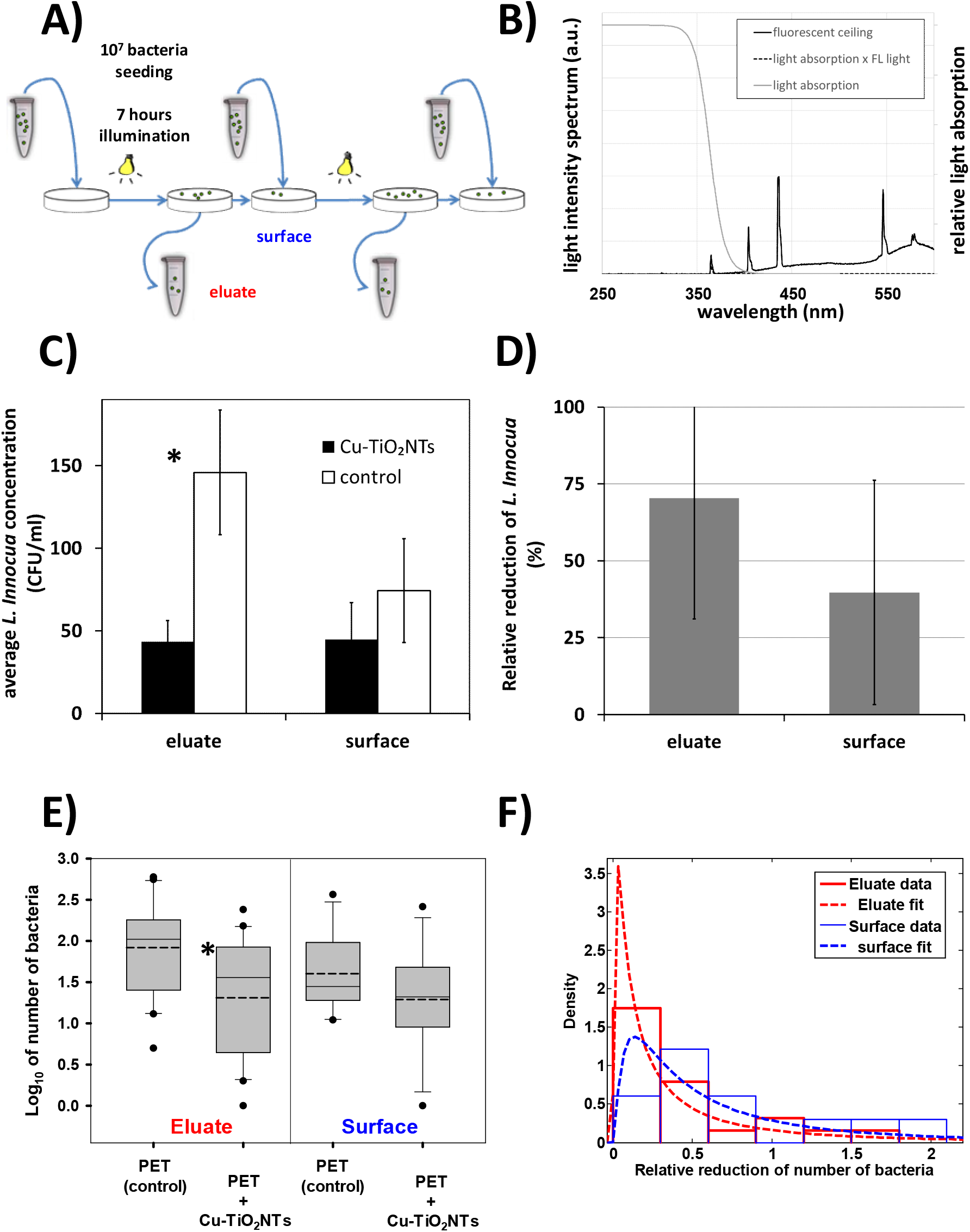
Survival of Listeria innocua under repeated daily contamination and washing. Reduction of number of bacteria Listeria innocua was measured on a polyethylene terephthalate (PET) surface with antibacterial nano coating (PET+Cu-TiO_2_NTs) or without the coating (PET) tested in a laboratory mimicking conditions in a food processing plant. A) Experimental setup scheme – Listeria innocua culture was continuously applied on surfaces as indicated by blue arrows pointing to a surface; B) spectrum of emited light from ceiling mounted fluorescent lights (black line); relative light absorption by the Cu-TiO_2_NTs (grey line); spectrum of absorbed light by the nanotubes (dashed line). C) Number of Listeria innocua shown on a logarithmic scale. On each surface 10^7^ bacteria were placed on PET slides. After 7 hours of exposure to ceiling mounted fluorescent lights remaining bacteria were transferred from the surfaces and colony forming units (CFU) were counted. Number of CFU on control surfaces is shown with white bars (PET control); Number of CFU on the nanoparticle coated surfaces is shown with black bars (PET + Cu-TiO_2_NTs); D) Relative reduction (%R) of bacteria as a consequnce of disinfecting action of nanoparticle coated surface, illuminated with ceiling mounted fluorescent lights (grey bars), calculated according to the equation 1 in Materials and Methods section. E) Number of Listeria innocua in eluate from either control surface without nano coating (PET) or from a surface coated with copper doped TiO_2_ nanotubes (PET+Cu-TiO_2_NTs) presented in a boxplot, where the line in a box marks the median number of bacteria, while the dashed line marks the mean. The boundary of the box closest to zero indicates 25^th^ percentile, and the boundary farthest from zero indicates the 75^th^ percentile. Whiskers (error bars) above and below the box indicate the 90^th^ and 10^th^ percentiles, respectively. The outliers are shown as dots. Note the LOG scale. Reduction in number of bacteria in eluate for 0.6 orders of magnitude was statistically significant^*^ (t = 3.018, with 38 degrees of freedom, two-tailed P-value = 0.00453, power of performed two-tailed test with alpha = 0.050: 0.837). The reduction of bacteria on a surface was smaller than in eluate (t = 1.338, with 20 degrees of freedom, two-tailed P-value = 0.196, power of performed two-tailed test with alpha = 0.050: 0.247). F) Histograms of the distribution (probability density function – PDF) of survival ratios in eluate (red) or on a surface (blue). Survival ratio was calculated as a ratio between the number of Listeria innocua from the surface with antibacterial coating (PET+Cu-TiO_2_NTs) and the number of Listeria innocua from a control surface without the coating (PET). For inefficient antibacterial coating survival ratio of 1 is expected. Dotted line is lognormal fit of the distributions with a maximum of the PDF at 0.02 and 0.13 for the reduction of the number of bacteria in eluate and on the surface, respectively.

Although the antimicrobial effect was not as pronounced as in the food processing plant, average number of CFU eluate from the control surface was 83 ± 4, which is significantly higher than the number of CFU in the eluate from the TiO_2_ nanotubes coated surface, which decreased to 21 ± 1 (*Figure 4* C, eluate) (t = 3.018, with 38 degrees of freedom, two-tailed P-value = 0.00453, power of performed two-tailed test with alpha = 0.050: 0.837). In the 28 days lasting experiment the average number of CFU remaining on the control surface was 40 ± 3, whereas the average number remaining on the nanotube coated surface decreased to 19 ± 4 (*Figure 4* C, surface). The relative reduction of the number of *Listeria Innocua* in the eluate was 70% ± 39 in seven hours of exposure to the fluorescent lights, compared to a control surface (*Figure 4* D, eluate). All the data, from which averages were calculated, are shown in in the supplement (Table S 1) and in the *Figure 4* E in a form of a box plot, from which can be easily seen that the distribution of measurements is not normal (the mean and median are not overlaping). For more detailed quantification of the results we therefore calculated also the ratios of bacteria from the coated surfaces versus the control surfaces for all measurements (*Figure 4* F). The effect of the antibacterial coating on the survival of *Listeria innocua* is indicated by the ratio of less than 1. The distributions of survival ratios were also not normal as for the data in *Figure 3* F. We again fit the histograms of the survival ratios in the eluate and on the surface. The best fits of the data to the lognormal distribution indicate that the maximum of the probability density function is around 0.1, thus confirming that antibacterial coating is inhibiting the growth of *Listeria innocua* on TiO_2_ nanotube coated surfaces.

To test whether a small addition of copper is solely responsible for the antibacterial properties of the nanotube coated surface, the above experiments were also performed in the dark (see the Supplement, Figure S 2). In the experiment in the dark the number of colony forming units of *Listeria innocua* on nanotube coated as well as on control surface was the same, indicating that the reduction of bacteria that we observed originates from photocatalytic process of the TiO_2_ nanotubes.

## Conclusion

To implement advantages of germicidal disinfection with the use of ultraviolet light as well as antibacterial properties of copper-containing surfaces, we used recently characterized Cu^2+^-doped TiO_2_ nanotubes and achieved a stable deposition on several materials, including on the surface of polyethylene terephthalate (PET), as the one of the synthetic polymers commonly used in food processing industry. More importantly, we showed that such coating has disinfecting effect, with the number of remaining microorganisms significantly decreased on the surface coated with Cu^2^+-doped TiO_2_ nanotubes as well as in eluate from the coated surface, when illuminated with common ceiling mounted fluorescent lights. The disinfection properties of the nanotube coated surfaces depend on the intensity of the light, which should include wavelengths at about 370 nm, as well as on the temperature difference of the surface and the dew point, where for maximum effectiveness of the photocatalytic effect the difference should be less than 2.5 °C.

Our results show that one dimensional nanomaterials, such as TiO_2_ nanotubes, can be employed for disinfection of polymer surfaces in the food industry, using cost effective illumination with existing fluorescent lights or additional low power light emitting diodes. Future use of such surfaces with antibacterial nano-coating and resulting sterilizing effect holds promise for such materials to be used in different environments or in better control of critical control points (HACCP) in food production as well as an improved biosecurity during the food manufacturing process.

## Acknowledgements

Special thanks to Maja Lepen for her excellent technical support in bacteriology laboratory. Work was funded by Slovenian Research Agency grant >Experimental biophysics of complex systems and imaging in biomedicine<, and by NAMASTE Centre of Excellence, Institute for research and development of Advanced Materials and Technologies for the Future.

## Contributions

J.Š., T.K., M.D., Š.P., and I.Z. conceived the experiments. P.U., T.K., Š.P., M.D., and I.Z. carried out the experiments. T.K., P.U., and Š.P. analyzed the data. J.Š., T.K., M.D., and Š.P. interpreted the results. M.D. and T.K. wrote the manuscript.

## Corresponding authors

Correspondence to prof. dr. Janez Štrancar and prof. dr. Martin Dobeic.

